# Examining motor evidence for the pause-then-cancel model of action-stopping: Insights from motor system physiology

**DOI:** 10.1101/2024.01.30.577976

**Authors:** Joshua R. Tatz, Madeline O. Carlson, Carson Lovig, Jan R. Wessel

## Abstract

Stopping initiated actions is fundamental to adaptive behavior. Longstanding, single-process accounts of action-stopping have been challenged by recent, two-process, ‘pause-then-cancel’ models. These models propose that action-stopping involves two inhibitory processes: 1) a fast Pause process, which broadly suppresses the motor system as the result of detecting any salient event, and 2) a slower Cancel process, which involves motor suppression specific to the cancelled action. A purported signature of the Pause process is global suppression, or the reduced corticospinal excitability (CSE) of task-unrelated effectors early on in action-stopping. However, unlike the Pause process, few (if any) motor system signatures of a Cancel process have been identified. Here, we used single- and paired-pulse TMS methods to comprehensively measure the local physiological excitation and inhibition of both responding and task-unrelated motor effector systems during action-stopping. Specifically, we measured CSE, short-interval intracortical inhibition (SICI), and the duration of the cortical silent period (CSP). Consistent with key predictions from the pause-then-cancel model, CSE measurements at the responding effector indicated that additional suppression was necessary to counteract Go-related increases in CSE during-action-stopping, particularly at later timepoints. Increases in SICI on Stop-signal trials did not differ across responding and non-responding effectors, or across timepoints. This suggests SICI as a potential source of global suppression. Increases in CSP duration on Stop-signal trials were more prominent at later timepoints. SICI and CSP duration therefore appeared most consistent with the Pause and Cancel processes, respectively. Our study provides further evidence from motor system physiology that multiple inhibitory processes influence action-stopping.

## Introduction

Our ever-changing environment sometimes requires us to quickly adjust our already-initiated actions. Action-stopping thus represents a fundamental component of flexible goal-directed behavior, and it is crucial to understand the underlying processes.

One widespread and longstanding view, proposed by Logan and Cowan (1984), holds that encountering a stop-signal triggers a control process that can inhibit an already initiated action. Whether successful action-stopping occurs ultimately hinges on whether this Stop process outpaces processes leading to action execution. This “horse-race” model views the Stop process as a single inhibitory process. By contrast, dual-process models, which include pause-then-cancel models of action-stopping (Diesburg & Wessel, 2021; Schmidt & Berke, 2017) propose two processes to underly action-stopping. First, the Pause process purportedly inhibits the motor system in response to any salient event, not just those requiring action-stopping. This inhibition is thought to be *global*, affecting both task-related and completely task-unrelated motor representations. Second, the Cancel process entails more focalized inhibitory control directed at the specific effector(s) needing stopping. Hence, a key prediction from these two-stage models is that motor representations targeted for action-stopping should show selective suppression (owing to the Cancel process) in addition to the initial non-selective suppression (owing to the Pause phase). The model also predicts that the Pause is stronger relatively early after the stop signal, whereas the Cancel process purportedly becomes stronger later. Fig. 1 depicts an example of these predictions.

**Figure 1.**
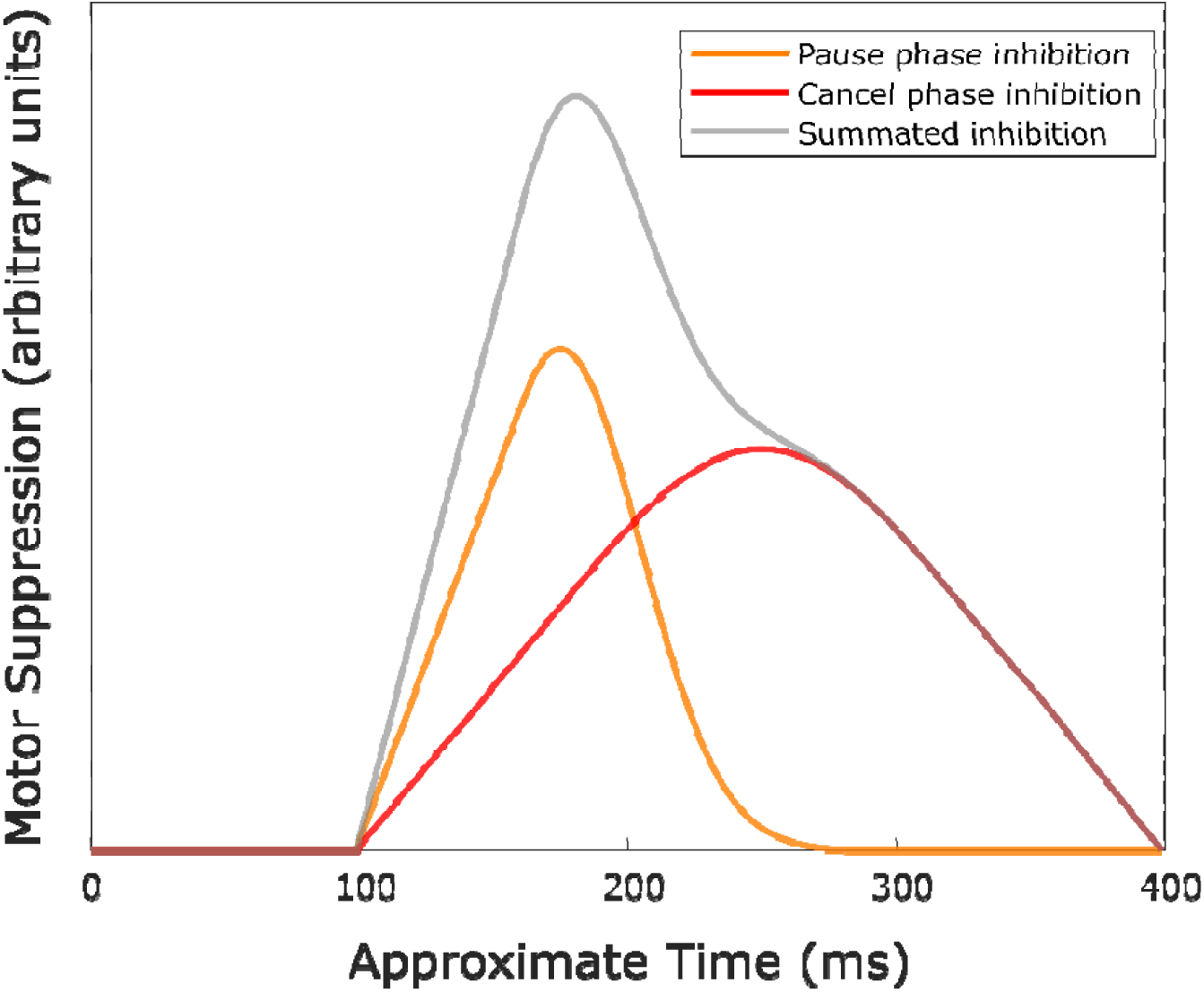
Example of motor suppression in a selected motor representation based on predictions from the pause-then-cancel model in humans (Diesburg & Wessel, 2021). The Pause process (Orange) peaks earlier than the Cancel process (Red). The result of summing the inhibition from each of these processes (Grey) is inhibition that is maximal at some intermediate timepoint. Note that, in this example, the Pause and Cancel processes onset at the same time; however, this need not be the case so long as the processes show some overlap in time. The Pause process is also assumed to be more transient (reflected as reduced variance above); however, the specific shape, amplitude, and timing of these distributions could vary from that above.

Evaluating these predictions necessitates neurophysiological measures with motor specificity and precise timing resolution. One of the most informative and physiologically meaningful measurements of action-stopping comes from motor evoked potentials (MEPs). MEPs are derived using single-pulse transcranial magnetic stimulation (TMS) and electromyography (EMG) and index the net corticospinal excitability (CSE) of specific muscles (e.g, Bestmann & Duque, 2016). During action-stopping, CSE is suppressed in both the specific muscle targeted for stopping and completely task-unrelated muscles (e.g., Badry et al., 2009; Cai et al., 2012; Wessel et al., 2013). This finding has often been linked to the horse-race model (e.g., Badry et al., 2009), and it has been argued that global inhibition could largely account for action-stopping ability (e.g., Raud et al., 2020). In other words, in a single-process view of action-stopping, this non-selective CSE suppression represents the entirety of the inhibitory effects exerted on the motor system during stopping.

However, a major challenge to this view is the finding that non-selective motor suppression is not actually unique to action-stopping. Indeed, salient events also induce non-selective CSE suppression, even in tasks that do not explicitly require stopping. Moreover, this suppression occurs at the same timepoints after these salient events and after stop-signals in action-stopping tasks (Dutra et al., 2018; Iacullo et al., 2020; Tatz et al., 2021; Wessel & Aron, 2013). On some successful stop trials, subthreshold EMG activity is observed in responding muscles that show a sharp downturn consistent with when non-selective CSE is observed (e.g., Jana et al., 2020; Raud & Huster, 2017; Raud et al., 2020). As with non-selective CSE, we found that subthreshold EMG was also found after salient events that required continuation of the Go response (Tatz et al., 2021).

While these neurophysiological signatures support the notion of a Pause process (Diesburg & Wessel, 2021), it is unclear whether similar neurophysiological measurements from motor cortex also suggest the presence of a Cancel process. In the current study, we measured CSE from both task-related and task-unrelated motor effectors. Participants alternated between blocks of manual and vocal stop-signal tasks (SST). CSE was measured from the right flexor digitorum superficialis (FDS), which was a primary agonist in the manual SST blocks, but completely task-unrelated in the vocal SST blocks (hereafter, we simply refer to CSE measurements from the task-unrelated effector as “non-selective CSE”). We measured CSE at timepoints where we expected maximal differences in the Pause and Cancel processes. Non-selective CSE suppression has been consistently found around 150 ms (cite) and has been proposed as a key signature of the Pause process (Diesburg & Wessel, 2021). The main proposed signature of the Cancel process is the frontocentral P3 response from EEG. This signature onsets around 200 ms and typically shows further amplitude increases around 250 ms (e.g., Wessel & Aron, 2015). We therefore measured CSE at 150 and 250 ms after the stop signal. Our main, pre-registered hypothesis was that selective and non-selective CSE suppression would peak at different time points after the stop-signal. Specifically, in line with the predictions from pause-then-cancel models, we hypothesized that non-selective CSE suppression would occur earlier than selective CSE suppression.

We also obtained two other TMS-derived measures that have been previously implicated in action-stopping to investigate whether either might show patterns consistent with Pause or Cancel processes. First, we used single- and paired-pulse TMS to measure short-interval cortical inhibition (SICI). Pharmacological studies have suggested that SICI reflects local GABA_A_ activity (e.g., Stagg et al., 2011). In bimanual stopping tasks, increased SICI has been found on successful stop trials in task-related effectors, including both the responding and non-responding hands (Chowdhury et al., 2019). As with CSE, the current study examined SICI from an effector when it represented a primary agonist and when it was completely task unrelated.

Second, we measured the cortical silent period (CSP), which is a time after the MEP offset during which the EMG trace shows no apparent activity (hence “silent”) prior to the resumption of normal EMG activity. CSP is considered to index the duration of local GABA_B_ activity (Werhahn et al., 1999). We analyzed CSPs during the manual task only. In general, few participants showed any CSPs in the vocal stopping task. Moreover, obtaining reliable CSPs in this task would have necessitated introducing isometric force requirements (e.g., Paci et al., 2021) that would have made the hand task related. During successful stopping, van den Wildenberg et al. (2010) demonstrated prolonged CSP durations shortly after the stop signal. The current work aimed to replicate and extend this work and test whether CSP aligns more closely with the purported timing of a Pause process (i.e., stronger effects early after the stop-signal) or a Cancel process (i.e., stronger effects later after the stop-signal).

## Methods

### Pre-registration and data availability

Pre-registration documents for Experiments 1 and 2 can be found on the Open Science Framework at https://osf.io/9mryu/ and at https://osf.io/rm7dz/. Data, task code, and analysis scripts will be uploaded to the same following manuscript acceptance after peer-review.

### Participants

Prior to participation, all participants completed a screening questionnaire including standard guidelines to determine TMS eligibility (Rossi et al., 2011) and gave their written informed consent. Additional eligibility criteria stipulated that participants must be right-handed, have self-reported normal vision, and be able to clearly articulate the letters “D”, “T”, “G”, and “K”. Participants for both experiments were recruited from Psychology courses, from the Iowa City community, and from among lab personnel. Participants either received course credit or were compensated at a rate of $15/hour. Upon full completion of the experiment, the entire meeting (including giving consent, hotspotting, etc.) lasted approximately 2-2.5 hours. The experiments were approved by the local Institutional Review Board (#201612707).

As detailed in our preregistration documents, we targeted an effective sample size of 25 participants for Experiment 1. This was determined based on the effective sample sizes from similar CSE studies (21-28 participants) from our group (Iacullo et al., 2020; Tatz et al., 2021). For Experiment 2, which began after the results of Experiment 1 were known, we targeted a sample size of 14 participants. This was determined with an *a priori* power analysis based on the large interaction effect found in Experiment 1 and assuming 1 group, 2 measurements, 90% power, and alpha = .05.

For Experiment 1 (single-pulse TMS only), 34 healthy adults were recruited. Data were excluded from 10 participants for various reasons (4 due to small or absent MEPs after motor hotspotting, 3 due to low trial counts in one or more conditions, 2 due to experimenter technical error, 1 due to an unanticipated computer update, and 1 due to an unanticipated fire alarm). This left an effective sample size of 24 participants (14 female, mean age = 20.12 years, SD age = 2.82 years) for Experiment 1. A sensitivity analysis in G*Power (Faul et al., 2009) with 80% power, alpha = .05, 1 group, and 6 repeated measures indicated power to detect at least medium sized effects (*f* = .22).

For Experiment 2 (single- and paired-pulse TMS), 19 healthy adults were recruited. None of the participants had participated in Experiment 1. Two of the participants in Experiment 2 were also experimenters, though these participants neither ran themselves nor processed their own data. Data were excluded from 5 participants due to various reasons (4 due to small or absent MEPs after motor hotspotting and 1 due to terminating the experiment early upon realization that TMS was producing headache and/or bothering scalp). This left an effective sample size of 14 participants (7 female, mean age = 21.93, SD age = 4.07) for Experiment 2.

### Manual and Vocal Stop-Signal Tasks

The participants alternated between completing manual and vocal versions of the SST. A task diagram is shown in Fig. 2. The tasks (and associated code) were adapted from prior work that examined non-selective CSE during SSTs for the voice (Wessel et al., 2022; 2016) and feet (Tatz et al., 2021). On every trial in the manual and vocal SSTs, participants made speeded responses to a go-signal consisting of a white letter centrally displayed on a black background. On 1/3 of trails, after some delay, the letter turned red, signifying the need to stop the response. This stop-signal delay (SSD) was initialized to 200 ms and adjusted adaptively in 50 ms increments (tracked separately for each response in each task). Participants had 1000 ms to indicate their response. The letter remained on the screen for the full response window. If no response was made within this time on a go trial, the message “Too Slow!” (in red) was displayed after the trial for .5 seconds. No message was displayed after stop trials. Inter-trial intervals of 3.2 and 3.7 s were used for Experiments 1 and 2, respectively. This allowed sufficient time for the TMS coil to recharge between trials.

**Figure 2.**
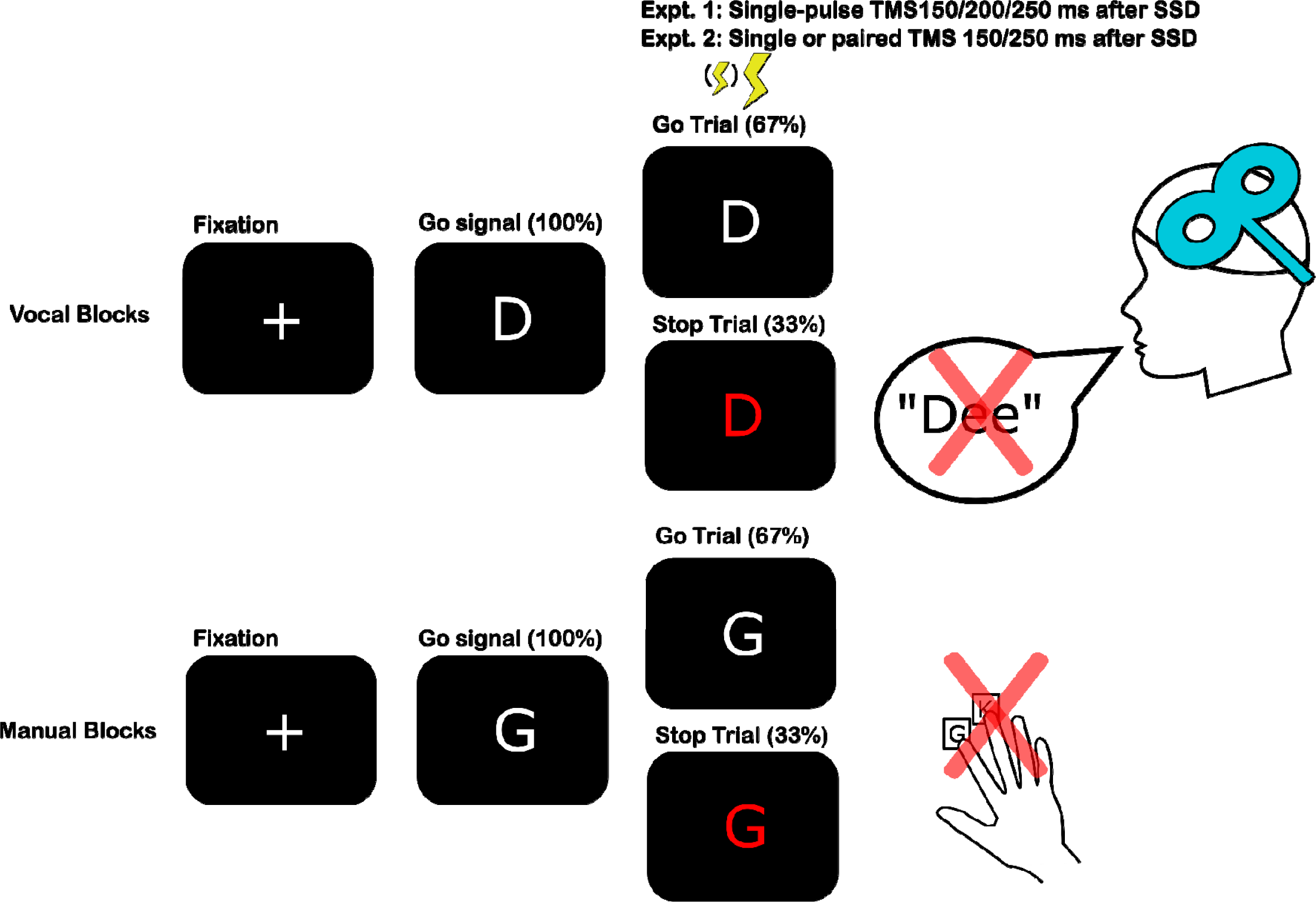
Illustration of the vocal and manual stop signal tasks as well as the timing and type of TMS stimulation for each experiment.

In the manual SST, participants made index and middle finger responses with their right hand to one of two letters (D and T or G and K) by depressing either the left arrow or the down arrow on a QWERTY keyboard (the arrow keys were selected so that participants fingers could rest in a comfortable position). In the vocal SST, participants read the displayed letter aloud.

Participants were instructed that their voice would need to be louder than the TMS click to be picked up by the microphone. Participants’ vocal responses were recorded by a USB-microphone for the duration of the response window (sampling rate = 48,000 Hz). The specific assignment of these letter pairings (D/T or G/K) to manual or vocal blocks were counterbalanced across two groups of participants. We also counterbalanced the order of these blocks such that one group of participants received manual blocks first (and on subsequent odd-numbered blocks) and the other group received vocal blocks first.

Most participants completed all 12 blocks which were each comprised of 90 trials (for a total of 1080 trials). However, three of the included participants in Experiment 1 and one participant in Experiment 2 completed 10 blocks.

### Voice data preprocessing

Voice data were preprocessed as in prior work from our group (Wessel et al., 2022; 2016). In brief, custom MATLAB scripts were used to playback the voice data from each trial and to visually inspect the audio data. Visualization of the audio data included information about the letter the participant was supposed to pronounce, whether the trial was a stop signal trial or not, and the algorithm-identified RT for that trial (if the amplitude threshold was crossed). The algorithm from prior work (Wessel et al., 2022; 2016) was designed to identify the RT as the first timepoint in which a set amplitude was exceeded (corresponding to the onset of the participant’s vocal response). However, the potential for a letter to be mispronounced or for the algorithm to misidentify the approximate onset of the vocal response required occasional editing. An analyst therefore inspected each voice trial and made adjustments if necessary. Typically, the RTs and Trial types were what the algorithm identified. On average, 3.31% of trials were edited in Experiment 1 and 4.67% in Experiment 2.

### EMG recordings

EMG data were recorded using adhesive electrodes (H124SG, Covidien) with belly-tendon placements. We selected the FDS of the right arm as a primary muscle involved in the manual task, and because MEPs from the FDS have been previously shown to not differ across index and middle finger responses in a choice RT task (Hasbroucq et al., 1997; 1999). One electrode of the bipolar pair was placed on the muscle belly (ventral forearm), and one was placed on the flexor tendons (near the wrist). An additional ground electrode was placed on the acromion process of the ipsilateral shoulder. The forearm was kept in a relaxed, pronated state during recordings. We also obtained EMG recordings from electrodes placed on the throat (with a corresponding ground electrode placed on the sternal end of the clavicle), although we did not analyze these data. EMG recording settings did not differ across Experiments 1 and 2. EMG recordings included the full trial duration or a matched time period during active baseline recordings.

### TMS stimulation settings, design, and procedure

For Experiment 1, TMS settings were similar to those reported in Tatz et al. (2021). The description is adapted from therein: TMS was delivered via a MagStim 200-2 system (MagStim) with a 70 mm figure-of-eight coil. Hotspotting was used to identify the correct location and intensity for each participant. We initially positioned the coil 5 cm left of and 2 cm anterior to the vertex and incrementally adjusted the location and intensity to determine the participant’s resting motor threshold. Resting motor threshold was defined as the minimum intensity needed to produce MEPs > 0.1 mV in 5 of 10 consecutive probes (Rossini et al., 1994). For Experiment 1, the stimulus intensity was increased to 115% of resting motor threshold. The mean experimental intensity was 63.1% of maximum stimulator output.

Most participants completed the full 1080 trials. Single-pulse TMS was delivered on 2/3 of those trials, with 360 pulses corresponding to stop trials and 360 to go trials. TMS was delivered 150, 200, or 250 ms after stimulus onset in equal numbers for each trial type. Thus, TMS was delivered on each stop trial. The remaining 360 trials were adjacent go trials that were dedicated to obtaining active baseline MEP measurements. Instead of delivering TMS during these trials, TMS was delivered during the inter-trial interval before the onset of the second go-signal on go trials. These recordings took place at 500, 700, or 900 ms after the termination of the preceding trial. Active baseline and no TMS trials were selected randomly from the list of all consecutive Go trials (with the constraint that equal numbers of 150, 200, and 250 ms stimulation times were preserved for regular go trials).

For Experiment 2, the TMS settings and procedure were the same as in Experiment 1, with some exceptions, given our interest in measuring SICI in this experiment. We therefore acquired both single- and paired-pulse TMS by changing the settings on the TMS device every two blocks from single-pulse mode to paired-pulse mode. On single-pulse blocks, we delivered a single pulse of TMS at 120% RMT, as the test pulse. On paired-pulse blocks, a conditioning pulse set at 80% RMT preceded the test pulse by 2 ms. These settings were selected based on previous resting-state SICI work from our group (Hynd et al., 2021). The mean test pulse was 62.1% and the mean condition pulse was 41.4% maximum stimulator output. The order in which participants received single- vs. paired-pulse blocks was counterbalanced across two groups of participants. Given the need to have single- and paired-pulse trials, we cut down on the number of trials per condition by limiting the test pulse stimulation times to 150 and 250 ms (i.e., excluding 200 ms stimulation time).

### TMS data preprocessing

The TMS data were preprocessed using custom MATLAB code that was modified from ezTMS, which is freely available MATLAB code for preprocessing TMS data with a graphical-user interface (Hynd et al., 2021). This enabled semi-automatic identification of TMS data characteristics and visual inspection of each trial. On each trial, an analyst who was not aware of the specific trial type (i.e., stop or go) visually inspected each trail and verified that the TMS pulse artifact and MEP were detectable on each trial with TMS stimulation. Identification of the TMS pulse artifact was facilitated by also visualizing the second voice EMG channel (which contained TMS pulse artifacts without corresponding MEPs). The MEP amplitude was extracted automatically and defined as the difference between the maximum and minimum deflection in the EMG trace 10-50 ms after the suprathreshold TMS pulse. Trials in which the MEP amplitude was less than .05 mV or in which the root mean square (RMS) of the EMG trace 80 ms before the TMS pulse exceeded .05 mV were automatically marked and rejected from all subsequent analyses. We also automatically rejected any trials in which the TMS pulse occurred at or prior to the corresponding RT for that trial. In Experiment 1, we also rejected the bottom 5% and top 5% MEPs in each condition. The decision to exclude these MEPs did not change any results. We did not do this for Experiment 2, given that there were slightly fewer potential MEPs per condition and that inclusion of the conditioning pulse in paired-pulse TMS would, by design, reduce MEP size.

As detailed in our pre-registration documents, participants’ data were included in the main analyses for Experiment 1 provided they had 15 usable MEPs for each Trial Type (Successful Stop, Go) in each Response Modality at each TMS Stim Time. In Experiment 2, this number was reduced to 10 usable MEPs.

### CSE

We calculated the mean amplitude of the non-rejected MEPs for each participant and each condition. The mean MEPs were then normalized with the appropriate active baseline MEPs (Voice or Hand). For Experiment 1, the average number of MEPs for each condition were as follows: 74.5 (Voice Baseline), 51.0 (Voice Go/150), 45.3 (Voice Go/200), 34.6 (Voice Go/250), 29.5 (Voice Successful Stop/150), 29.4 (Voice Successful Stop/200), 28.2 (Voice Successful Stop/250), 77.1 (Hand Baseline), 50.75 (Hand Go/150), 43.5 (Hand Go/200), 34.5 (Hand Go/250), 26.4 (Hand Successful Stop/150), 30.5 (Hand Successful Stop/200), 30.8 (Hand Successful Stop/250).

For Experiment 2, these numbers on single-pulse trials were: 44.6 (Voice Baseline), 39.9 (Voice Go/150), 30.2 (Voice Go/250), 22.2 (Voice Successful Stop/150), 22.4 (Voice Successful Stop/250), 44.7 (Hand Baseline), 38.1 (Hand Go/150), 28.4 (Hand Go/250), 22.2 (Hand Successful Stop/150), 24.1 (Hand Successful Stop/250). And these numbers for paired-pulse trials were: 41.9 (Voice Baseline), 36.6 (Voice Go/150), 26.6 (Voice Go/250), 20.1 (Voice Successful Stop/150), 19.4 (Voice Successful Stop/250), 42.6 (Hand Baseline), 37.4 (Hand Go/150), 26.9 (Hand Go/250), 19.1 (Hand Successful Stop/150), 23.4 (Hand Successful Stop/250).

### SICI

Experiment 2 enabled measurement of SICI, which was calculated according to the following formula:

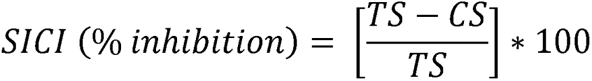

Where TS refers to the suprathreshold (120% RMT) test stimulus and CS refers to the subthreshold (80% RMT) condition stimulus. Here, SICI is framed as % inhibition (e.g., Wadsley et al., 2023) with greater positive values indicating greater inhibition (i.e., more SICI).

### CSP

Mean CSP (absolute cortical silent period) on Hand trials and the corresponding active baseline was examined in both experiments. During the CSP, EMG activity is suppressed even below baseline. For each trial with an MEP, an analyst naïve to the specific Trial Type (stop, go) visually determined whether a CSP was present and, if so, manually marked the timepoint where normal EMG activity resumed. An algorithm then refined the time point to be the maximum point within +/- 2 ms of the selected point. In general, CSPs were not observed on Voice trials, when the motor representation targeted with TMS was unrelated to the task. By contrast, CSPs were present on a preponderance of Hand trials, including the active Hand baseline trials. Only CSPs resulting from single-pulse TMS that produced valid MEPs were included in the CSP analyses.

### Data analyses

Data were analyzed in JASP 0.18.1 (Love et al., 2019) using RM-ANOVAs (with Hunyh-Feldt correction where applicable) and Bayesian RM-ANOVAs (with non-informative priors). Positive findings from these analyses were followed-up on with simpler ANOVA models or paired *t*-tests (with Holm-Bonferroni correction in the case of frequentist comparisons). Bayes factor is reported as the strength of evidence for the null (*BF_01_*) or alternative (*BF_10_*) hypothesis according to the following convention: inconclusive (*BF* ∼ 1), anecdotal (*BF* > 1), moderate (*BF* > 3), and strong (*BF* > 10).

For the behavioral data, mean RTs on Go and failed stop trials were submitted to RM-ANOVAs that included Trial Type (Go, Failed Stop) and Response Modality (Manual, Vocal) as factors. Errors and misses were generally rare, and thus not analyzed. Stop-signal reaction time (SSRT) was calculated via the integration method (Verbruggen et al., 2019). Paired *t*-tests were used to compare SSRT across the manual and vocal SSTs.

Of main interest were the TMS data on successful Stop compared to Go trials. The CSE data from Experiment 1 were analyzed according to the 2 (Trial type: Successful Stop, Go) x 2 (Response Modality: Manual, Vocal) x 3 (TMS Stim Time: 150, 200, 250) within-subject design. The CSE and SICI data from Experiment 2 were analyzed in the same fashion, albeit, without the 200 ms TMS Stimulation Time. The CSP data in both experiments were analyzed with the within-subject factors Trial Type and TMS Stim Time.

Of secondary interest, we completed similar analyses to those above but this time comparing successful and failed stop trials. Generally, more failed stop trials were excluded because increased EMG was detected just before the TMS pulse and RT more often occurred before the TMS pulse. This was expected since failed stop trials represent the faster portion of the go RT distribution. However, our requirement to have at least 10 MEPs for each failed stop condition for CSE analyses constrained the analyses that we could do. In Experiment 1, all 24 participants had over 10 usable failed stop trials per task (Manual, Vocal) at the 150 ms Stim Time, and 18 of these participants met this inclusion criteria at the 200 ms Stim Time. However, only four participants met this inclusion criteria for 250 ms. Our CSE analyses with failed stop trials were thus limited to the 18 participants who presented at least 10 usable failed stop trials at the 150 and 200 ms TMS Stim Times. Given that Experiment 2 likewise showed substantial dropout at the late stimulation time and did not include an intermediate stimulation time as Experiment 1 had, we refrained from analyzing CSE and SICI on failed stop trials in this experiment. For the CSP data, we decided to include data so long as at least five usable trials were available from each failed stop condition. This allowed us to compare failed stop CSPs from 20 participants from across the two experiments.

Lastly, we examined to what extent the pause-then-cancel model vs. other possibilities could account for data at the individual-subject level. For each participant, each Response Modality, and each TMS Stim Time, we calculated Cohen’s d based on the difference in CSE on Go and successful stop trials. For each Cohen’s d in the manual task, we additionally subtracted from this value the Cohen’s d at the corresponding timepoint in the vocal task. Thus, the effect sizes in the manual task reflected additional CSE suppression beyond the observed non-selective CSE at the same time. For each participant we first identified whether they showed evidence of non-selective CSE suppression and/or selective CSE suppression. To be considered as such, at least one of the timepoints had to have a Cohen’s d > .2 (corresponding to a small effect). For participants who met the criteria for showing both non-selective and selective CSE suppression, we determined the order by identifying the non-selective and selective timepoint showing the greatest increase in CSE suppression compared to previous timepoints. To be considered an increase, the increase in Cohen’s d had to be > .2. Each participant was then classified as global-then-selective, selective-then-global, same time global and selective, global only, selective only, or neither.

## Results

### Behavior

The RT results for both experiments are displayed in Fig. 3. The RM-ANOVA on go and failed stop RTs in Experiment 1 indicated a main effect of Trial Type, *F*(1, 23) = 467.63 , *p* < .001, 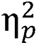 = .95, *BF_10_* = 7.69x10^11^, demonstrating that mean RTs were faster on failed stop trials than on go trials. This relationship held within each SST, as well (Manual: *t*(23) = 21.06, *p* < .001, *d* = 4.30, *BF_10_* = 3.09x10^13^; Vocal: *t*(23) = 12.00, *p* < .001, *d* = 2.45, *BF_10_* = 4.01x10^8^). There was also a main effect of Response Modality, *F*(1, 23) = 28.16, *p* < .001, 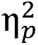 = .55, *BF_10_* = 837, and a significant Trial Type X Response Modality interaction, *F*(1, 23) = 46.18, *p* < .001, 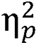 = .67, *BF_10_* = 2.55x10^5^. These indicated that RTs were generally faster for vocal responses than hand responses, and that these differences were more pronounced on failed stop trials (simple effect: *F*(1, 23) = 53.02, *p* < .001, 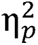 = .70, *BF_10_* = 3.20x10^5^) than on go trials (simple effect: *F*(1, 23) = 12.71, *p* = .002, 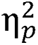 = .36, *BF_10_* = 29.7). SSRTs did not significantly differ across tasks, and there was moderate Bayesian evidence to support this lack of difference, *t*(23) = .07, *p* = .943, *d* = .015, *BF_01_* = 4.65. The mean SSD and mean p(inhibit) for the manual SST were, respectively: 265 ms (SD = 69 ms) and .506 (SD = .034). For the vocal SST, these numbers were: 221 ms (SD = 53 ms) and .520 (SD = .080).

**Figure 3.**
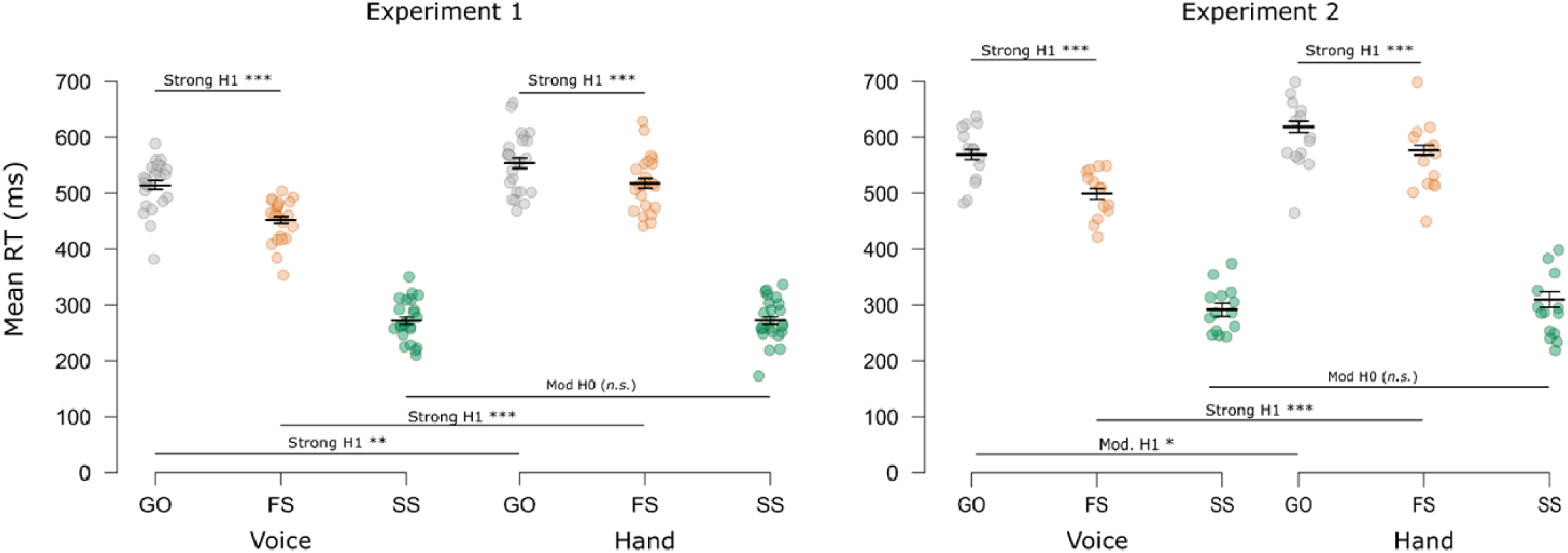
Mean behavioral RTs for Go and failed stop (FS) trials and mean SSRTs (SS). Gray and orange dots indicate the individual mean RTs for Go and FS trials, respectively. Green dots indicate mean SSRTs (calculated via the integration method, Verbruggen et al., 2019). Short horizontal bars indicate the group mean RT/SSRT. Error bars indicate SEM. Long horizontal bars indicate multiple comparison testing. Strong H1/H0: BF > 10; Mod. H1/H0: BF > 3; Anec. H1/H0: BF > 1.3; H1: alternative; H0: null hypothesis. ***p < 0.001; **p < 0.01; *p < 0.05; n.s., p > .05.

The results for Experiment 2 mirrored those of Experiment 1. We again observed main effects of Trial Type, *F*(1, 13) = 234.43, *p* < .001, 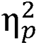 = .95, *BF_10_* = 4.11x10^5^; Response Modality, *F*(1, 13) = 16.77, *p* = .001, 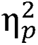 = .56, *BF_10_* = 34.3; and the Trail Type x Response Modality interaction, *F*(1, 13) = 14.13, *p* = .002, = .52, *BF_10_* = 42.55. Importantly, failed stop RTs were faster than go RTs in both SSTs (Manual: *t*(13) = 7.96, *p* < .001, *d* = 2.13, *BF_10_* = 8017; Vocal: *t*(13) = 13.23, *p* < .001, *d* = 3.54, *BF_10_* = 1.67x10^6^). Vocal responses were again faster than manual responses and more so on failed stop trials (simple effect: *F*(1, 13) = 24.57, *p* < .001, = .65, *BF_10_* = 146) than on go trials (simple effect: *F*(1, 13) =7.73, *p* = .016, = .37, *BF_10_* = 3.52). SSRTs did not significantly differ across tasks, which was again supported by moderate Bayesian evidence, *t*(13) = 0.26, *p* = .797, *d* = 0.07, *BF_01_* = 3.60. The mean SSD and mean p(inhibit) for the manual SST were, respectively: 302 ms (SD = 75 ms) and .494 (SD = .035). For the vocal SST, these numbers were: 262 ms (SD = 60 ms) and .511 (SD = .010).

### CSE

The main CSE results for both experiments are shown in Fig. 4. Corresponding plots depicting the CSE differences (go minus successful stop) are displayed in Fig. 5.

**Figure 4.**
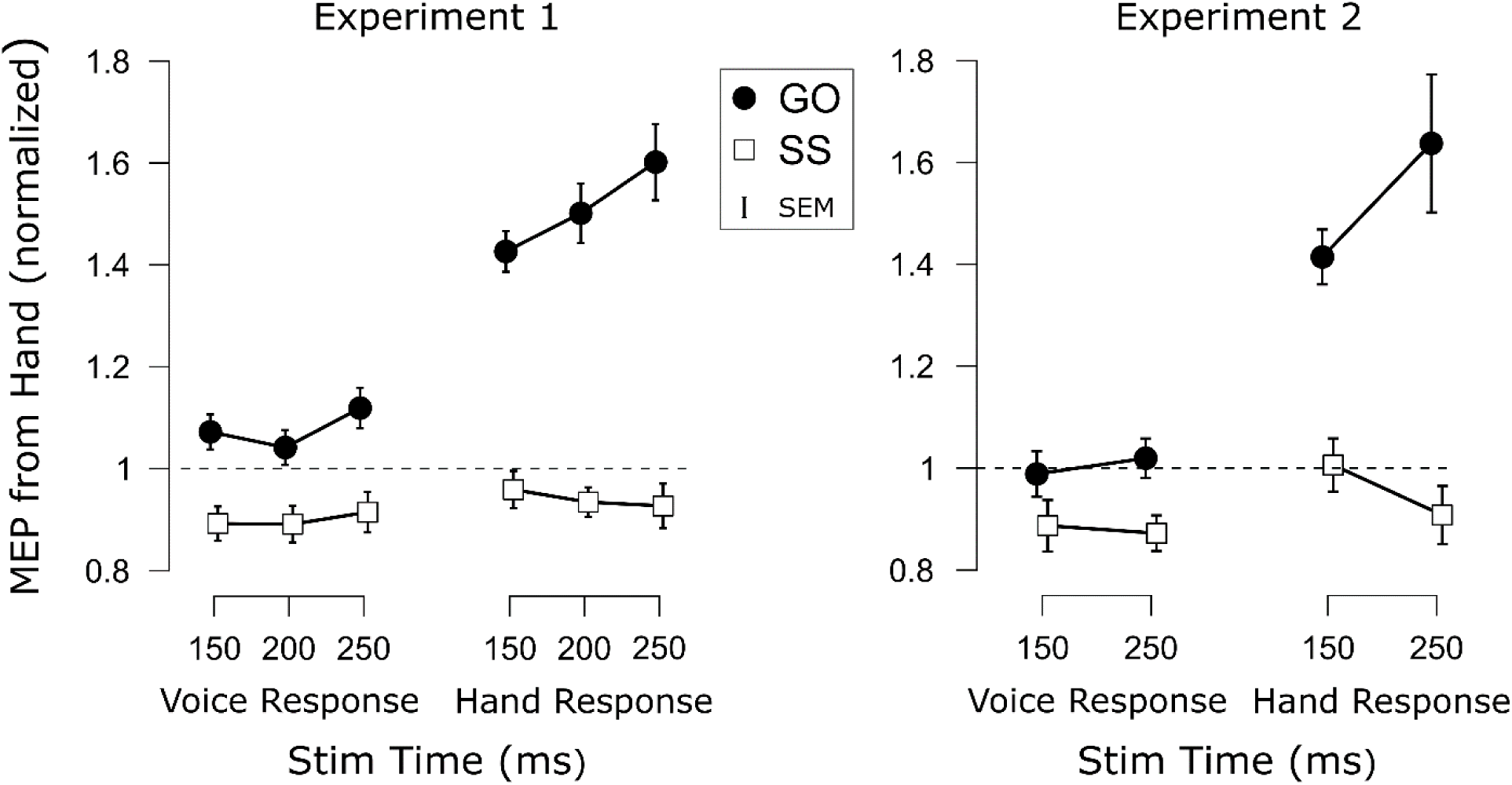
Mean baseline normalized corticospinal excitability (CSE) recorded from the flexor digitorum superficialis for each Trial Type (Go, SS = Successful Stop), Task (Voice, Hand), and TMS Stimulation Time in each experiment. Points indicate the group mean for each condition. Error bars indicate SEM.

**Figure 5.**
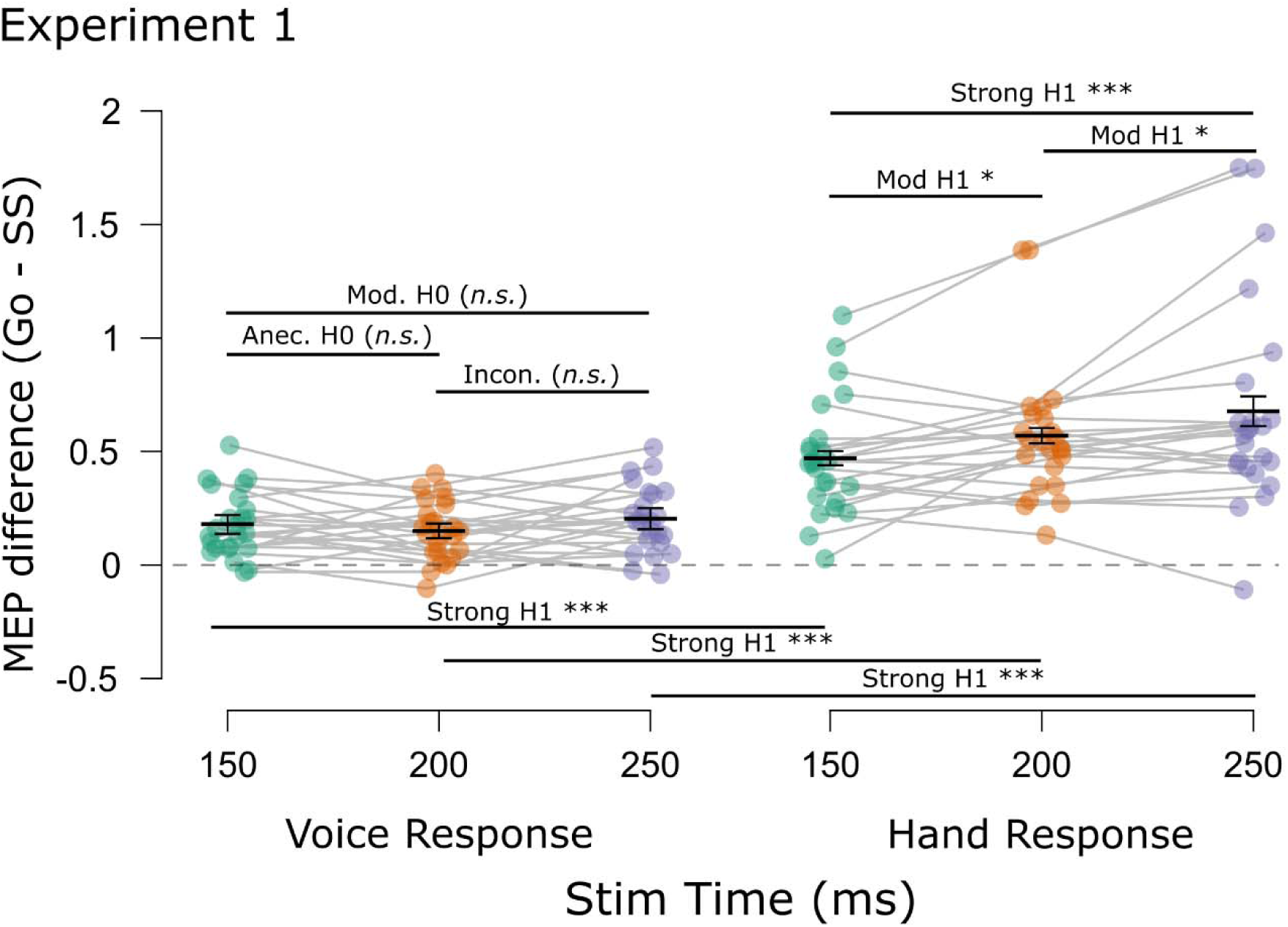
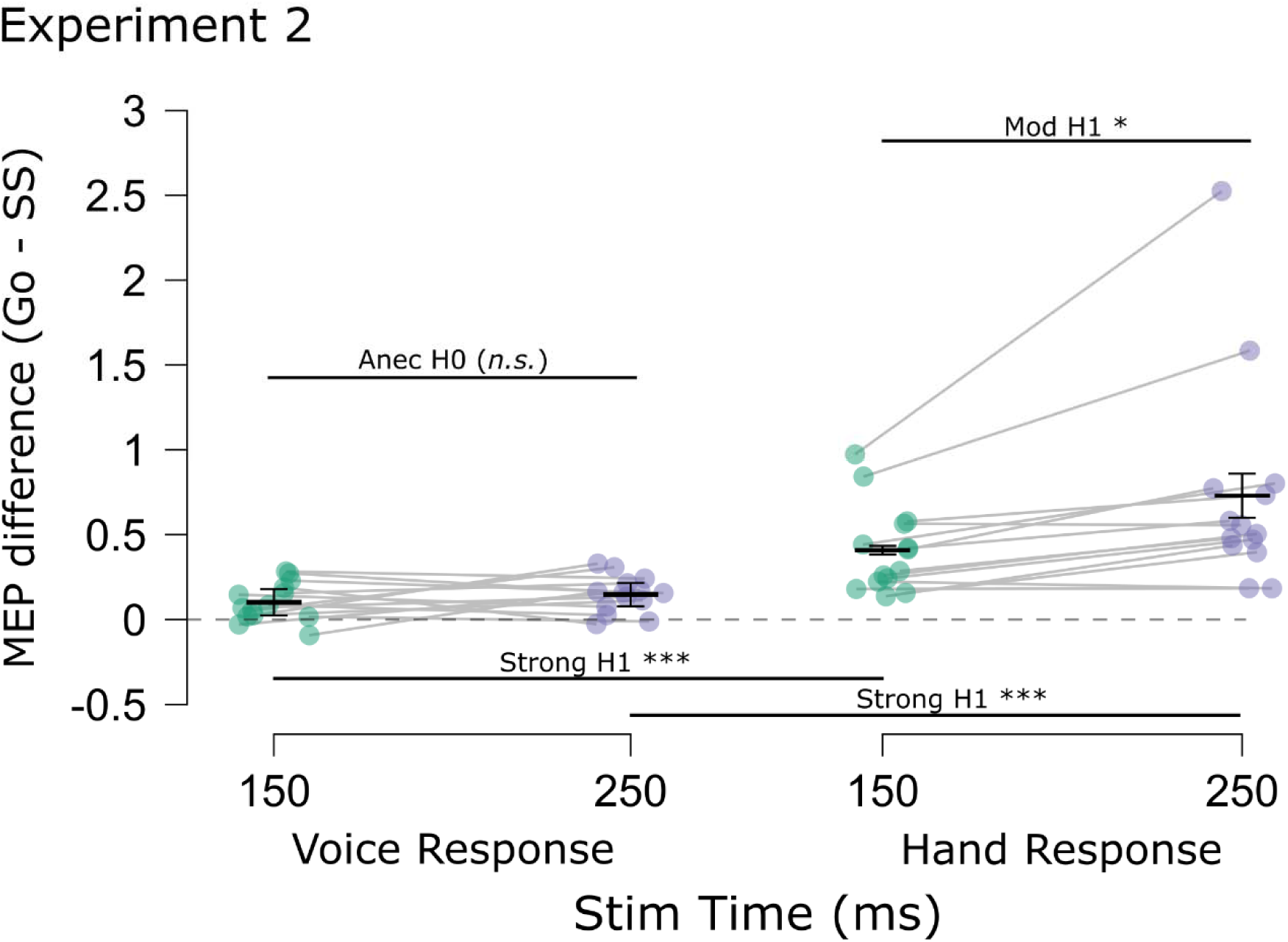
Mean difference in CSE (Go minus successful Stop) for each Task (voice, hand) and TMS Stimulation Time in each experiment. Points indicate mean CSE differences for individual subjects, Light gray horizontal bars connect means from the same subjects. Short black horizontal bars indicate the group mean CSE differences. Error bars indicate SEM. Long black horizontal bars indicate multiple comparison testing. Strong H1/H0: BF > 10; Mod. H1/H0: BF > 3; Anec. H1/H0: BF > 1.3; Incon. H1/H0: BF ∼ 1. H1: alternative; H0: null hypothesis. ***p < 0.001; *p < 0.05; n.s., p > .05.

Following our predictions derived from the pause-then-cancel model in humans (Diesburg & Wessel, 2021), of main interest was whether we would detect selective CSE suppression that was additional to non-selective CSE suppression. If so, we expected this additional selective CSE suppression to occur at later time points. First, there was a main effect of Trial Type, *F*(1, 23) = 97.64, *p* < .001, 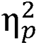 = .81, *BF_10_* = 1.22x10^7^, consistent with considerable prior work showing that CSE is suppressed on successful stop trials (e.g., Badry et al., 2009; Coxon et al., 2006). There were also main effects of Response Modality, *F*(1, 23) = 14.43, *p* < .001, 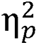 = .37, *BF_10_* = 29.3; and of TMS Stim Time, *F*(1.83, 42.08) = 9.69, *p* < .001, 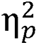 = .30, *BF_0_* = 10.36. Also significant were the two-way Trial Type x Response Modality, *F*(1, 23) = 36.69, *p* < .001, 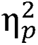 = .62, *BF_10_* = 4895; and the Trial Type x TMS Stim Time interactions, *F*(1.65, 37.97) = 8.52, *p* = .002, 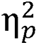 = .27, *BF_10_* = 24.2; but not the Response Modality x TMS Stim Time interaction, *F*(1.76, 40.37) = 0.89, *p* = .448, 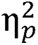 = .03, *BF_01_* = 5.85. Most importantly, though, the three-way Trial Type x Response Modality x TMS Stim Time was significant, *F*(1.70, 38.99) = 5.20, *p* = .013, 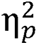 = .18, *BF_10_* = 33.1. To interpret this interaction with respect to our question of main interest, it is most useful to compare the degree of CSE suppression at each timepoint (see Fig. 5). To this end, we compared CSE suppression at the same timepoint across Response modalities and across timepoints within each Response Modality (9 comparisons). Compared to non-selective CSE suppression, greater selective CSE suppression was found at each TMS Stim Time (150 ms: *t*(23) = 5.17, *p_Holm_* < .001, *d* = 1.06, *BF_10_* = 1582; 200 ms: *t*(23) = 7.36, *p_Holm_* < .001, *d* = 1.50, *BF_10_* = 1.80x10^5^; 250 ms: *t*(23) = 4.81, *p_Holm_* < .001, *d* = .98, *BF_10_* = 704). Comparing CSE suppression across timepoints within each Response Modality yielded further insight. Non-selective CSE suppression did not significantly differ across any timepoints (150 vs. 200: *t*(23) = 1.09, *p_Holm_* = .289, *d* = .22, *BF_01_* = 2.75; 200 vs. 250: *t*(23) = 2.03, *p_Holm_* = .054, *d* = .41, *BF_10_* = 1.22; 150 vs 250: *t*(23) = 0.76, *p_Holm_* = .46, *d* = .16, *BF_01_* = 3.58). Moreover, Bayes factor provided anecdotal evidence for no difference at 150 vs. 200 ms and moderate evidence for no difference at 150 vs. 250 ms. Although the evidence for 200 vs. 250 ms was inconclusive, reflecting numerically greater non-selective CSE at 250 ms, it is worth noting that the non-selective CSE at 250 ms did not differ from that at 150 ms. In contrast to non-selective CSE suppression, the selective CSE suppression increased with each successive timepoint (150 vs. 200: *t*(23) = 2.41, *p_Holm_* = .012, *d* = .49, *BF_10_* = 4.57; 200 vs. 250: *t*(23) = 2.38, *p_Holm_* = .013, *d* = .49, *BF_10_* = 4.31; 150 vs 250: *t*(23) = 3.61, *p_Holm_* = .004, *d* = .74, *BF_10_* = 49.8). Thus, the Experiment 1 results were consistent with the notion that additional selective CSE suppression was increasingly necessary at later timepoints to counteract the excitatory influences associated with the Go process.

In general, the Experiment 2 data showed the same basic patterns as Experiment 1 (Fig. 4). There was a main effect of Trial Type, *F*(1, 13) = 38.43, *p* < .001, = .75, *BF_10_*= 886, indicating successful stop trials generally showed suppressed CSE relative to go trials. And again, there was a main effect of Response Modality, *F*(1, 13) = 19.50, *p* < .001, = .60, *BF_10_* = 51.0. No main effect of TMS Stim Time was found this time, *F*(1, 13) = 0.98, *p* = .341, = .07, *BF_01_*= 2.46. More importantly, though, the same two- and three-way interactions were found as in Experiment 1 (Trial Type x Response Modality, *F*(1, 13) = 13.58, *p* = .003, = .51, *BF_10_* = 16.01; Trial Type x TMS Stim Time: *F*(1, 13) = 8.39, *p* = .013, 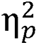 = .39, *BF_10_* = 3.37; Response Modality x TMS Stim Time: *F*(1, 13) = 0.98, *p* = .341, 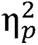 = .07, *BF_01_* = 2.27; Trial Type x Response Modality x TMS Stim Time: *F*(1, 13) = 6.92, *p* = .021, 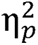 = .35, *BF_10_* = 7.92). As in Experiment 1, we completed multiple comparisons (4 comparisons) on the CSE suppression differences (Fig. 5). As before, greater selective CSE suppression was found at each timepoint compared to non-selective CSE suppression (150 ms: *t*(13) = 3.91, *p_Holm_* = .006, *d* = 1.05, *BF_10_* = 24.0; 250 ms: *t*(13) = 3.45, *p_Holm_* = .004, *d* = .92, *BF_10_* = 11.4). Moreover, whereas non-selective CSE suppression did not differ across timepoints, *t*(13) = 1.07, *p_Holm_* = .304, *d* = .286, *BF_01_* = 2.28, selective CSE suppression was greater at 250 ms compared to 150 ms, *t*(13) = 2.97, *p_Holm_* = .011, *d* = 0.79, *BF_10_* = 5.31. Thus, the Experiment 2 CSE findings joined those of Experiment 1 in suggesting that additional selective CSE suppression emerged at later timepoints.

As planned in our pre-registration, we also took an individual differences approach to examining the CSE data. Based on separately computed effect sizes for each Response Modality and TMS Stim Time, we classified each participant as global-then-selective, selective-then-global, same time global and selective, global only, selective only, or neither. For Experiment 1 (with three TMS Stim Times), we identified 10 participants as global-then-selective, 8 as selective-then-global, 4 as same time global and selective, 1 as global only, 1 as selective only, and 0 as neither. For Experiment 2 (with two TMS Stim Times), we identified 6 as global-then-selective, 5 as selective-then-global, 1 as same time global and selective, 1 as global only, 1 as selective only, and 0 as neither. Thus, while we expected a majority of participants to be classified as global-then-selective, this classification held only a slight majority (42.1% of participants) over the next most common selective-then-global group (34.2%). Nevertheless, these results clearly indicate that additional selective suppression is apparent at the individual level, as very few participants showed global only or selective only suppression.

We now consider the results from the failed stop trials in Experiment 1 from the subset of participants who presented enough usable failed stop trials at the 150 and 200 ms TMS Stim Times. These results are displayed in Fig. 6. We found significant main effects of Trial Type, *F*(1, 17) = 55.68, *p* < .001, 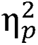 = .77, *BF_10_* = 1.57x10^4^; of Response Modality, *F*(1, 17) = 18.78, *p* < .001, 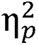 = .53, *BF_10_* = 70.0; and of TMS Stim Time, *F*(1, 17) = 8.57, *p* = .009, 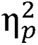 = .34, *BF_10_* = 2.48 (anecdotal evidence). Also significant were the two-way interactions of Trial Type x Response Modality, *F*(1, 17) = 31.43, *p* < .001, 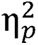 = .65, *BF_10_* = 902; and of Trial Type x TMS Stim Time, *F*(1, 17) = 6.77, *p* = .019, 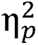 = .29, *BF_10_* = 1.91 (anecdotal evidence). However, the Response Modality x TMS Stim Time interaction was not significant, *F*(1, 17) = 3.14, *p* = .094, 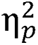 = .16, *BF_01_* = 1.19 (inconclusive). Finally, the Trial Type x Response Modality x TMS Stim Time was nearly significant and was supported by near moderate Bayesian evidence, *F*(1, 17) = 4.10, *p* = .059, 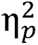 = .19, *BF_10_* = 2.86. As with our comparisons of successful stop and go trials, we followed-up on these analyses by comparing the failed stop minus successful stop differences in CSE across tasks and timepoints. Although CSE was (at least numerically) elevated on all failed stop trials relative to stop trials, these differences were much larger with selective CSE (150 ms: *t*(17) = 4.82, *p_Holm_* < .001, *d* = 1.14, *BF_10_ =* 186; 200 ms: *t*(17) = 4.90, *p_Holm_* < .001, *d* = 1.15, *BF_10_ =* 215). Differences were not found across timepoints for non-selective CSE, *t*(17) = 0.98, *p_Holm_* = .340, *d* = .23, *BF_01_* = 2.70 (anecdotal). Thus, this echoed our findings involving go and successful stop differences, although it may be worth noting that the failed stop trials here occurred at levels intermediate to those on go and successful stop trials (suggesting failed stop trials prompted some non-selective CSE suppression). By contrast, selective CSE showed larger differences at 150 ms than at 200 ms, *t*(17) = 2.47, *p_Holm_* = .050, *d* = .58, *BF_01_* = 2.54 (anecdotal). Interestingly, this is the opposite of the pattern what was seen with go and successful stop differences. This suggests that additional selective CSE suppression did not kick in until later timepoints on failed stop trials.

**Figure 6.**
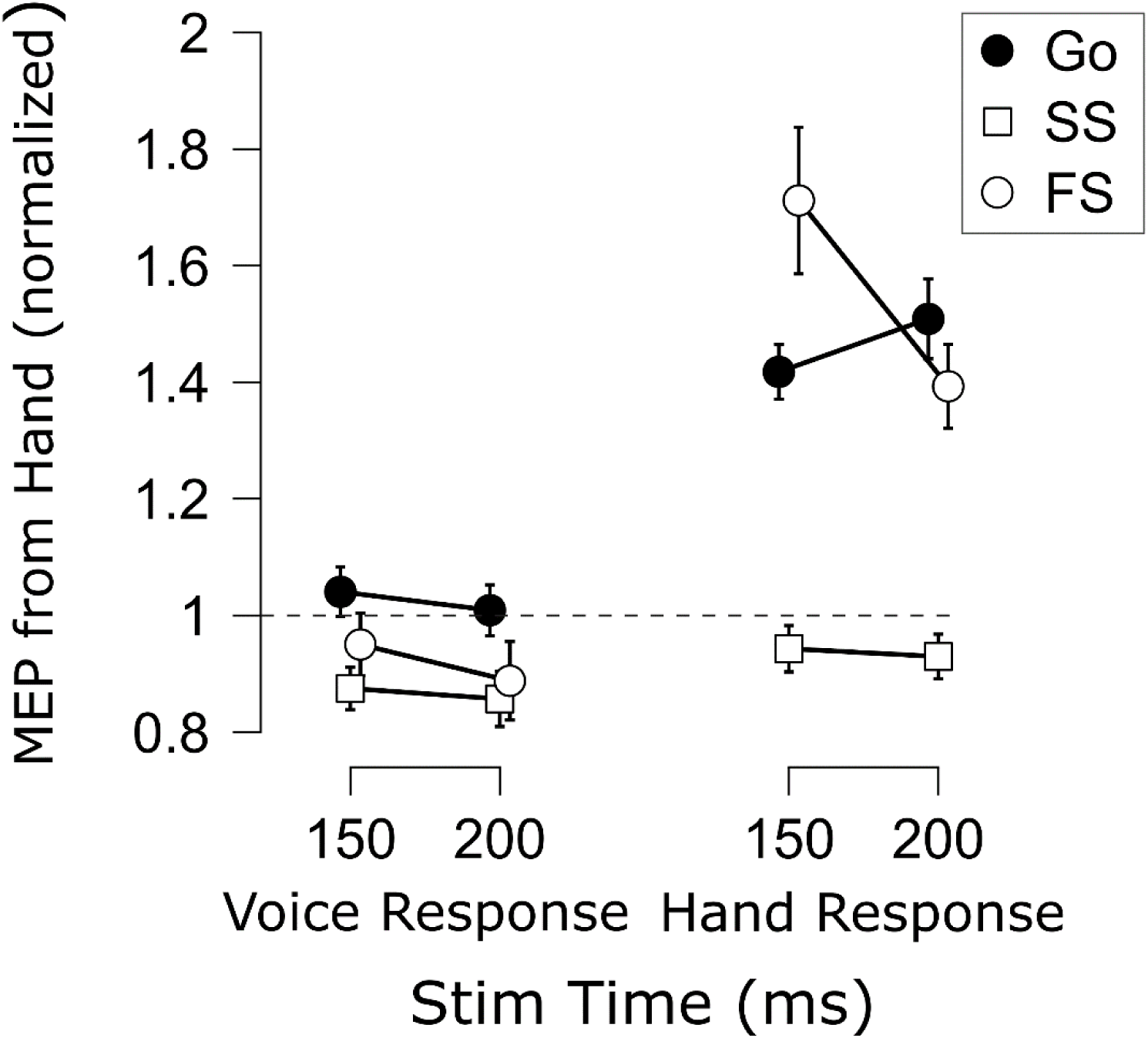
Mean baseline normalized corticospinal excitability (CSE) for each Trial Type (GO, SS = Successful Stop, FS = Failed Stop), Task (Voice, Hand), and TMS Stimulation from 18 participants in Experiment 1 who provided FS MEPs in sufficient numbers (≥ 10 MEPs per condition). Points indicate group means. Error bars indicate SEM.

### SICI

The SICI results are displayed in Fig. 7. In line with prior work (e.g., Chowdhury et al., 2019; Coxon et al., 2006), there was a main effect of Trial Type, *F*(1, 13) = 8.47, *p* = .012, = .40, *BF_10_* = 4.96, indicating that SICI was generally elevated on successful stop compared to go trials. There was also a main effect of Response Modality, *F*(1, 13) = 88.36, *p* < .001, = .87, *BF_10_* = 5.57x10^4^, indicating that SICI was increased when the probed motor representation was non-selected compared to when the same representation was selected. Interestingly, these SICI differences were apparent even during the active baseline. There was no main effect of TMS Stim Time on SICI, *F*(1, 13) = 0.05, *p* = .824, = .00, *BF_01_* = 4.00. The two- and three-way interactions were also not significant (Trial Type x Response Modality: *F*(1, 13) = .038, *p* = .849, = .00, *BF_01_* = 2.64; Trial Type x TMS Stim Time: *F*(1, 1. 13) = .142, *p* = .712, = .01, *BF_01_*= 2.93; Response Modality x TMS Stim Time: *F*(1, 13) = 3.01, *p* = .11, = .19, *BF_10_* = 1.31; Trial Type x Response Modality x TMS Stim Time: *F*(1, 13) = .128, *p* = .726, = .01, *BF_01_* = 6.76). Of course, it may be worth noting that evidence for the Response Modality x Time interaction was inconclusive. This seemed to reflect the at least numeric trend by which non-selective SICI seemed to increase over time, whereas selective SICI seemed to decrease over time. In the Discussion section, we will point out why we think this might be an interesting avenue to pursue in future research. For the current focus on examining SICI with respect to the human pause-then-cancel model of action-stopping, it is important to note that even if these seeming increases and decreases in SICI over time are interpreted as such, they were present to similar degrees on both successful stop and go trials. This is corroborated by examining the % differences in SICI (stop minus go) shown in Fig. 8. Indeed, these comparisons indicated near-moderate to moderate Bayesian evidence for a lack of differences in stop-related SICI inhibition across tasks and TMS Stim Time (Vocal/150 ms vs. Vocal/250 ms: *t*(13) = 0.73, *p* = .477, *d* = .20, *BF_01_* = 2.94; Manual/150 ms vs. Manual/250 ms: *t*(13) = 0.10, *p* = .920, *d* = .03, *BF_01_* = 3.69; Vocal/150 ms vs. Manual/150 ms: *t*(13) = 0.04, *p* = .967, *d* = .01, *BF_01_* = 3.70; Vocal/250 ms vs. Manual/250 ms: *t*(13) = 0.27, *p* = .788, *d* = .07, *BF_01_* = 3.58). Thus, the stop-related SICI increases appeared to be sustained over time and with global consequences to the motor system.

**Figure 7.**
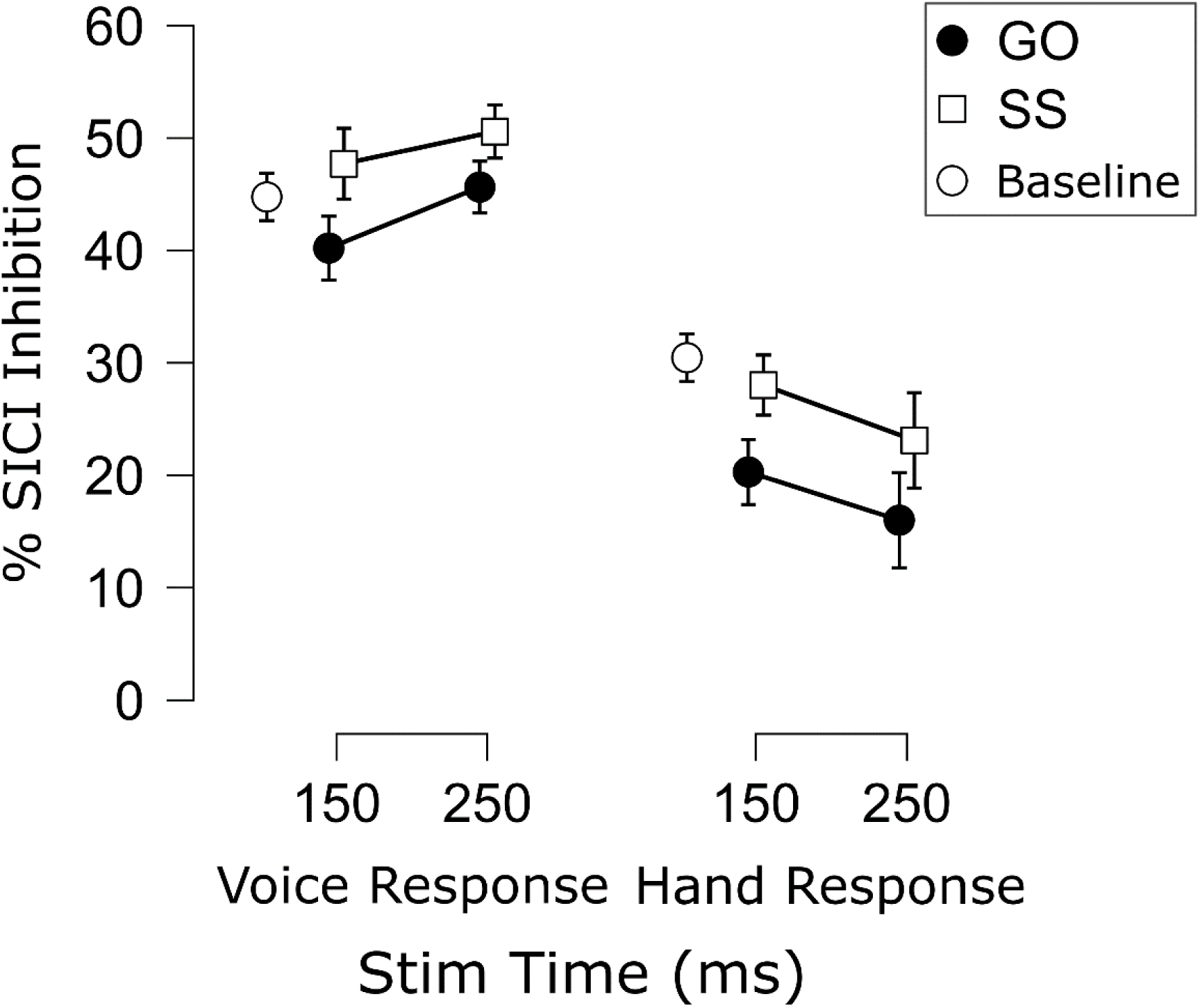
Mean % SICI inhibition [(Test stimulus – Conditioning stimulus)/Test stimulus * 100] for each Trial Type (Go, SS = Successful Stop), Task (Voice, Hand), and TMS Stimulation Time in Experiment 2. Points indicate the group mean for each condition. White circles indicate group mean % SICI inhibition during the active baseline in each Task. Error bars indicate SEM.

**Figure 8.**
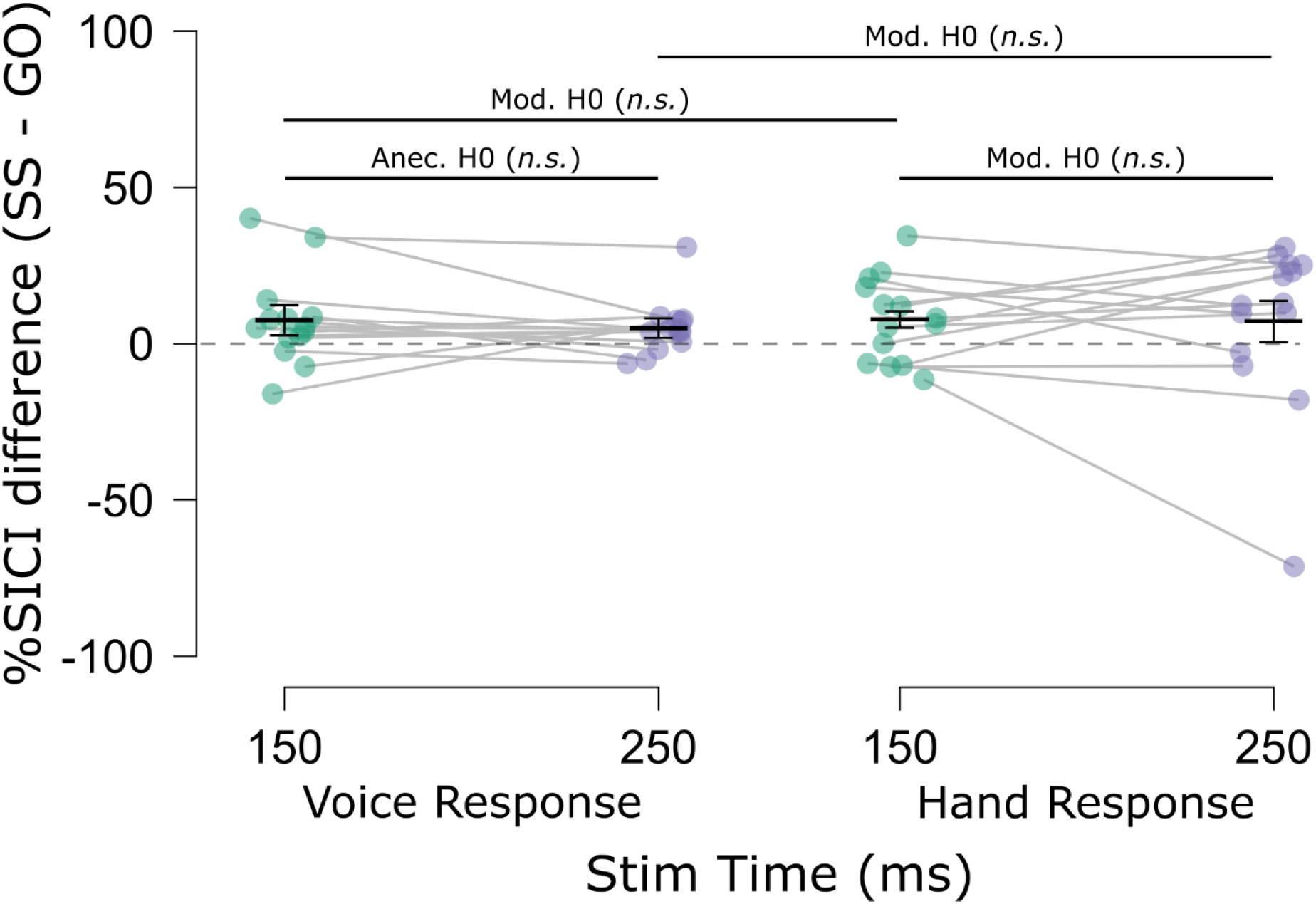
Mean % SICI inhibition differences (Successful stop minus Go) for each Task (voice, hand) and TMS Stimulation Time in Experiment 2. Points indicate mean % SICI differences for individual subjects, Light gray horizontal bars connect means from the same subjects. Short black horizontal bars indicate the group mean % SICI differences. Error bars indicate SEM. Long black horizontal bars indicate multiple comparison testing. Strong H1/H0: BF > 10; Mod. H1/H0: BF > 3; Anec. H1/H0: BF > 1.3; H1: alternative; H0: null hypothesis. ***p < 0.001; *p < 0.05; n.s., p > .05.

### CSP

The CSP data from each experiment are displayed in Fig. 9. In Experiment 1, there was a main effect of Trial Type in which CSPs were longer on successful stop trials compared to go trials, *F*(1, 20) = 16.47, *p* < .001, 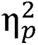 = .45, *BF_10_* = 59.2. It is worth noting that this increase in CSP duration on successful stop trials brought the mean CSP duration on these trials closer to, but still below the duration observed at baseline. No main effect of TMS Stim Time was found, *F*(1, 20) = 1.11, *p* = .341, 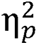 = .05, *BF_01_* = 6.45. More importantly, however, there was a significant interaction of Trial Type x TMS Stim Time, *F*(1.77, 35.40) = 6.37, *p* = .006, 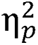 = .24, *BF_10_* = 46.2. To interpret the interaction, follow-up paired *t*-tests (6 comparisons) were performed to assess potential differences in successful stop and go trials at each stimulation time and whether any of these differences differed across stimulation times. Although, the numerical difference in successful stop and go CSP durations did not reach significance and was inconclusive at 150 ms, *t*(20) = 1.92, *p*_Holm_ = .069, *d* = .42, *BF_10_* = 1.06, CSPs were clearly prolonged on successful stop trials at the later timepoints (200 ms: *t*(20) = 1.92, *p*_Holm_ = .069, *d* = .42, *BF_10_* = 51.7; 250 ms: *t*(20) = 1.92, *p*_Holm_ = .069, *d* = 0.42, *BF_10_* = 103). When comparing these CSP duration differences (stop minus go) across TMS Stim Times, the difference at 250 ms was clearly greater than at 150 ms, *t*(20) = 2.99, *p*_Holm_ = .029, *d* = 0.65, *BF_10_* = 6.60. There was also a greater difference at 200 compared to 150 ms, although the *p*-value did not survive correction for multiple comparisons and the Bayesian evidence was near moderate, *t*(20) = 2.53, *p*_Holm_ = .060, *d* = 0.55, *BF_10_* = 2.86. Moderate evidence for no difference was found between 200 and 250 ms, *t*(20) = 0.65, *p*_Holm_ = .52, *d* = 0.14, *BF_01_* = 3.62. We therefore interpret the interaction as indicating that CSP differences between successful stop and go increased at later timepoints.

**Figure 9.**
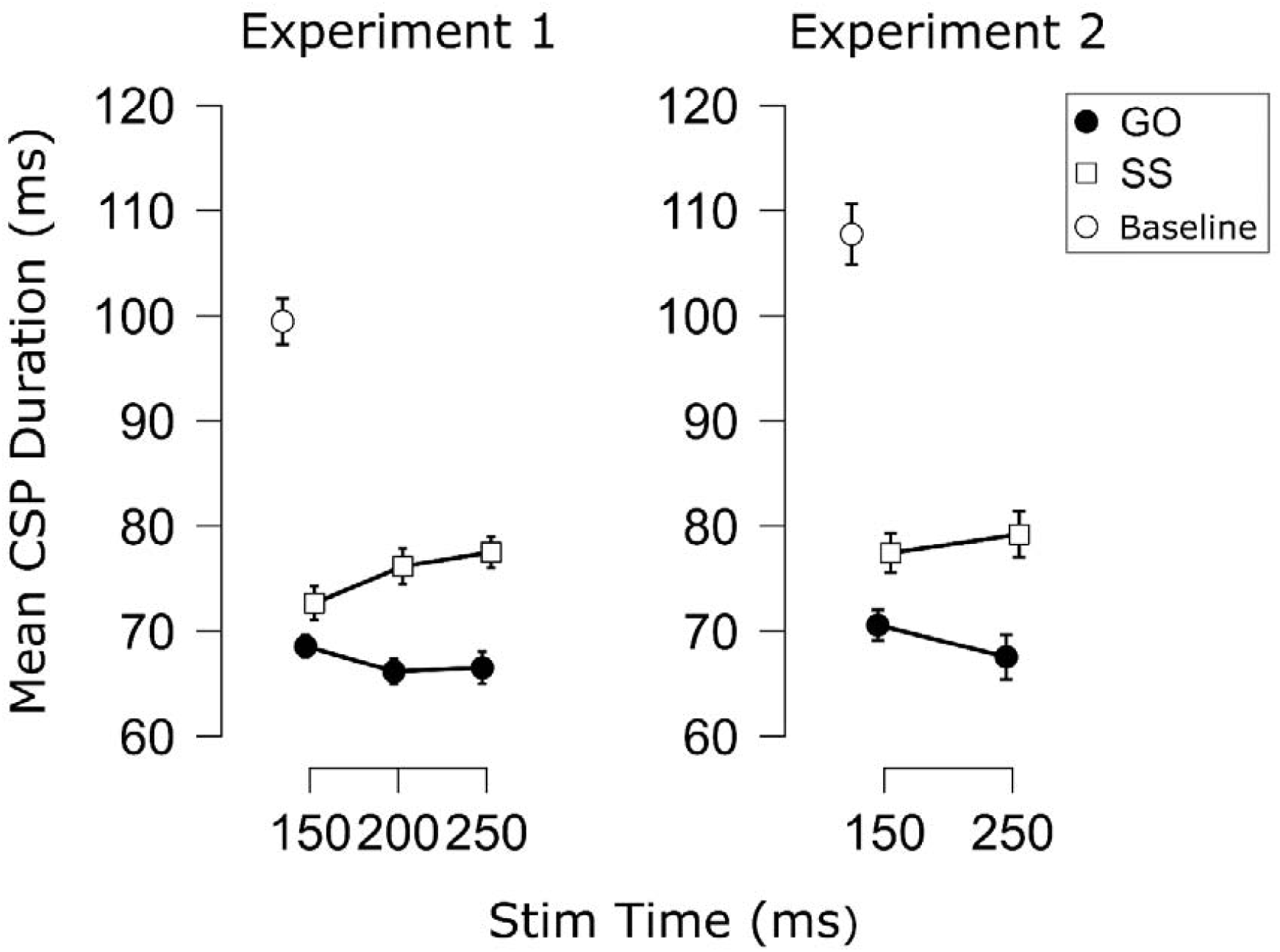
Mean cortical silent period (CSP) duration (in ms) for each Trial Type (Go, SS = Successful Stop) and TMS Stimulation Time during the manual SST in each experiment. Points indicate the group mean for each condition. White circles indicate the group mean CSP duration during the baseline measurement between trials. Error bars indicate SEM.

The CSP results from Experiment 2 (Fig. 9), showed similar patterns to those in Experiment 1. Again, a main effect of Trial type was found indicating decreased CSPs on successful stop trials, *F*(1, 13) = 15.95, *p* = .002, = .55, *BF_10_* = 4.50. Again, no main effect of time was found, *F*(1, 13) = 0.09, *p* = .765, = .00, *BF_01_*= 2.67. This time, the interaction did not reach significance and the Bayesian evidence (at least with non-informative priors) was anecdotal, *F*(1, 13) = 3.55, *p* = .082, = .21, *BF_10_* = 1.33. However, it is worth noting that the CSP difference still increased numerically across stimulation times and the corresponding effect size was similar in magnitude to that of Experiment 1 ( of .21 compared to .24). We therefore think the reduced power of this experiment to be the most likely explanation for the lack of statistically significant effect, here.

As with the CSE data, there were considerably fewer subjects with sufficient failed stop CSPs at late timepoints. We therefore combined the CSP data across Experiments 1 and 2, as this enabled comparing failed stop, successful stop, and go trials with 20 participants at both early and late timepoints (150 and 250 ms). The data are shown in Fig. 10. Again, a main effect of Trial Type was found, *F*(1.62, 30.72) = 10.96, *p* < .001, 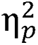 = .37, *BF_10_* = 155, but no main effect of TMS Stim Time, *F*(1, 19) = 1.75, *p* = .202, 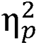 = .08, *BF_01_* = 3.58. More importantly, the Trial Type X TMS Stim Time interaction was significant, *F*(1.70, 32.34) = 4.42, *p* = .025, 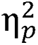 = .19, *BF_10_* = 6.12. Posthoc comparisons were then used to compare the CSP duration on failed stop trials to the other Trial Types at each time point. There did not appear to be a difference in CSPs durations between failed stop trials and go trials at 150 ms, *t*(19) = 1.12, *p*_Holm_ = .279, *d* = .25, *BF_01_* = 2.49 (anecdotal); but CSP duration did seem to be reduced compared to successful stop trials, *t*(19) = 2.53, *p*_Holm_ = .021 (uncorrected), *d* = .57, *BF_10_* = 2.82 (anecdotal-near moderate). At 250 ms, the CSP duration on failed stop trials was increased compared to go trials, *t*(19) = 2.92, *p*_Holm_ = .036, *d* = .65, *BF_10_*= 5.70; however, this CSP duration was still reduced compared to successful stop trials, *t*(19) = 2.59, *p*_Holm_ = .018 (uncorrected), *d* = .58, *BF_10_* = 3.17. This would suggest that the same process that drives CSP prolongation on successful stop trials at later timepoints is also active on failed stop trials. However, as failed stop trials showed decreased CSP duration (numerically below go) even at earlier timepoints, it would seem plausible that initial variation in CSP duration at early timepoints may account for action-stopping outcomes.

**Figure 10.**
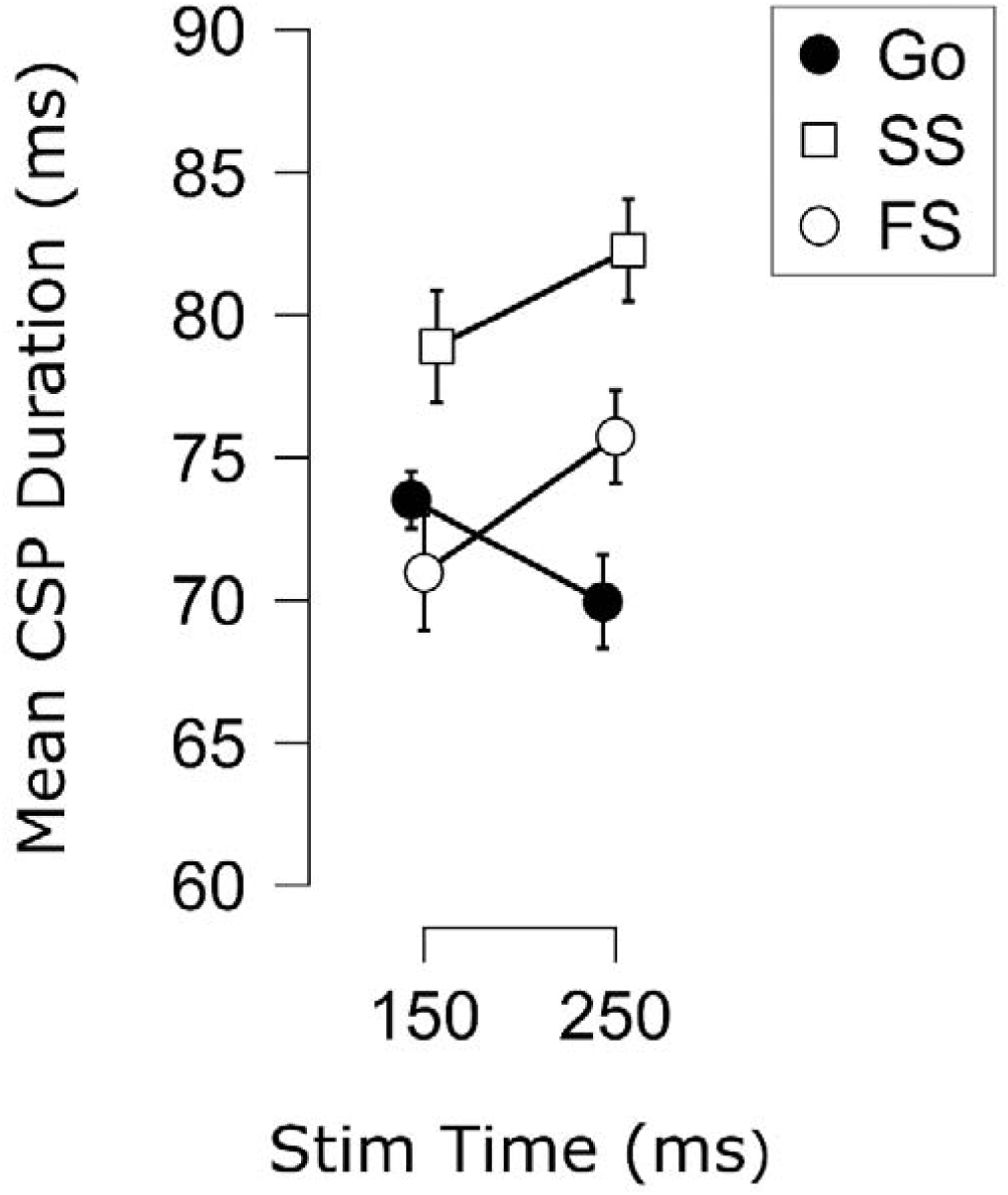
Mean cortical silent period (CSP) duration (in ms) for each Trial Type (Go, SS = Successful Stop, FS = Failed Stop) and TMS Stimulation Time from the 20 participants across the two experiments with FS CSPs in sufficient numbers (> 5 CSPs per condition). Points indicate the group mean for each condition. Error bars indicate SEM.

## Discussion

The current study measured CSE, SICI, and CSP to probe motor activity during action-stopping. Importantly, we systematically manipulated the task-relevance of the same muscle (FDS) to isolate non-selective and selective sources of suppression at early and late stimulation times. This allowed us to assess predictions of the pause-then-cancel model in humans (Diesburg & Wessel, 2021) vis-à-vis single-process models.

### CSE evidence for Pause and Cancel processes

As our main means of examining evidence for the Cancel process, we used single-pulse TMS to measure CSE selectively and non-selectively. CSE has the advantage of capturing net excitability, or the sum of excitatory and inhibitory influences, for a given motor representation (e.g., Bestmann & Duque, 2016). This is important because inhibitory control need not be achieved by local inhibitory neurons. For instance, inhibitory control could be achieved by truncated input, and this is rather what is proposed to be the case in both the Cancel process (Diesburg & Wessel, 2021) and in some single-process models, such as the interactive-race model (Boucher et al., 2007). Thus, we considered CSE to be ideal for evaluating our main prediction from the pause-then-cancel model, which was that task-related suppression should peak *later* than task-unrelated suppression.

Our CSE data from two experiments suggest that this is the case. Specifically, we found that selective CSE on successful stop trials was decreased compared to go trials, and this was especially the case at later timepoints, after non-selective suppression had already peaked. Failed stop trials likewise showed decreased CSE from 150 to 200 ms. As matched go timepoints showed CSE increases at this time, this further suggests the emergence of a unique stopping process sometime after 150 ms. We therefore consider these results to be more consistent with a two-stage, pause-then-cancel model.

Some CSE results were unexpected, though. First, owing to the notion of the Pause process as an early, more-transient process, we expected that non-selective CSE suppression would likely wane around 250 ms. Instead, the group-level CSE results indicated that non-selective CSE suppression was roughly equivalent across all stimulation timepoints (i.e., 150, 200, and 250 ms). However, even if the Pause process is less transient than thought, what is more important for pause-then-cancel models is that the selective suppression showed a separate time course that was later-emerging than the non-selective suppression. Second, we expected a clear majority of participants to show a pattern of global-then-selective suppression (as we found at the group-level). However, a similar number showed selective-then-global suppression. Future work is necessary to examine whether these represent stable subgroups, or if these results reflect the limited timepoints we sampled. Nevertheless, the individual-level CSE results indicate both non-selective and selective sources of suppression, as very few participants showed only one type of suppression.

### Evidence for global effects of SICI

As with CSE, we examined SICI from an effector when it was task related and when it was completely task unrelated. Prior work has shown increased SICI on successful stop trials compared to go trials (e.g., Chowdhury et al., 2019; Coxon et al., 2006). However, it remained an open question whether increased SICI during stopping would be found in a completely task unrelated effector (Derosiere & Duque, 2020).

More importantly for pause-then-cancel model considerations, the increases in SICI we found on successful stop trials did not differ across timepoints or based on the task-relevance of the effector. Subsequently, successful stopping would seem to be characterized by global increases in SICI and may, at least in part, account for the increases in non-selective CSE suppression during successful action-stopping. Indeed, we found SICI to be increased above baseline (at least numerically speaking) when the effector was not selected, which matches the below baseline non-selective (global) CSE suppression commonly seen during action-stopping (e.g., Badry et al., 2009) and in the current study. Also, we did not find stop-related SICI to change over time, just as was found with non-selective CSE. Within the pause-then-cancel model, our SICI findings were therefore most consistent with a Pause process, even if, as with non-selective CSE, the stop-related increases in SICI were less transient than expected. Aside from having global effects on the motor system, the other main Pause process prediction is that the inhibition be triggered by any salient event. Compared to go signals in an anticipatory response inhibition task, Wadsley et al. (2023) showed similar increases (at least numerically) in SICI in a task-relevant effector after both stop-signals and salient signals instructing response continuation.

Taken with our finding of global effects of SICI, these studies provide initial evidence for SICI as a potential signature of the Pause process.

Other, ostensibly non-stopping related features of our SICI data were also noteworthy. First, compared to non-selective SICI, selective CSE was much-reduced even during the active baseline period. This would seem in line with delayed-response tasks in which decreased SICI is found in task-related effectors (especially the selected effector) in the delay period (Duque & Ivry, 2009). Second, although inconclusive in terms of Bayes factor (and non-significant), we observed an at least numeric trend where, if anything, SICI decreased over time on both stop and go trials when selected (in the Manual task) and increased over time when non-selected (in the Vocal task). Although we do not wish to overinterpret an inconclusive finding, the direction of these effects are at least in line with Derosiere and Duque’s (2020) review in which SICI has been shown to decrease during action selection while increasing in other, non-selected effectors (which were either still task-related or anatomically related to the currently selected action). In future work, it may be interesting to consider whether SICI increases related to action selection extend to even completely unrelated effectors.

### Action-stopping marked by prolonged CSPs, particularly at late timepoints

Across experiments, we found increased CSP on successful stop trials, with small (inconclusive/non-significant) differences at 150 ms becoming much larger at later timepoints (200 and 250 ms). Failed stop trials showed a similar magnitude increase in CSP duration from 150 to 250 ms, as well, despite starting at a level comparable to go trials. This suggests the emergence of a unique stop-related process occurring sometime after 150 ms. Notably, these findings paralleled our selective CSE results, and match the proposed timing characteristics of the Cancel process, which should at least peak at these later timepoints.

Whereas CSP duration on stop trials increased back toward baseline over time, CSP duration on go trials sustained or showed further decreases. This suggests that CSP duration may reflect a tonic inhibitory state that is lifted during the Go process but counteracted when action-stopping is required. This view would seem consistent with the notion that the Cancel process involves shutting off excitatory drive associated with the Go process (Diesburg & Wessel, 2021).

Our findings are also largely consistent with van den Wildenberg et al. (2010), who showed increased CSP duration on successful stop trials beginning around 134 ms after the stop signal with further increases around 179 ms. Of course, these differences are earlier than what we observed and then what we would expect for a Cancel phase signature. Aside from different stimulation times, one notable difference between our studies was that van den Wildenberg et al. used a simple-response SST whereas we used a choice-response SST. Potentially, this could have sped up inhibitory processes in their task. It is also worth noting that the early increase in CSP duration they found occurred at a time when the CSP duration on go trials showed a decreasing trend. We think this early difference could be explained by an early, non-selective source of suppression, potentially reflecting the Pause process. Notably, Rangel et al. (2023) recently tasked participants with completing an SST with the legs while maintaining isometric force in the hand to measure motor output continuously and non-selectively. They found isometric force to be significantly reduced beginning 183 ms post-signal onset (among earlier numeric differences). Potentially, van den Wildenberg et al.’s use of isometric force to measure CSP with TMS might have enabled the detection of earlier differences than would have been possible with isometric EMG alone. Future work could assess this possibility by combining the current work with van den Wildenberg et al. (2010) and Rangel et al. (2023) to examine CSP duration both selectively and non-selectively. This could inform the interesting possibility of whether early CSP prolongation reflects the Pause process and late CSP prolongation reflects the Cancel process. While a limitation of the present work is that we did not do this and thus cannot gauge potential non-selective contributions to CSP duration, we think it most important that we replicated van den Wildenberg et al. (2010) in finding stop-related increases in CSP that increased over time. Regardless of which specific processes are at play, our findings join van den Wildenberg et al. in implicating CSP prolongation as a signature of action-stopping.

In summary, we examined CSE, SICI, and CSP from a motor representation both when it was involved in an action-stopping task and when it was completely unrelated to the task. The timing and motor-selectivity of our CSE results were consistent with predictions from the recently proposed pause-then-cancel model in humans (Diesburg & Wessel, 2021). Consideration of more advanced TMS measures provided additional insights. SICI (like non-selective CSE suppression) had global effects on the motor system. CSP duration (like selective CSE suppression) peaked at relatively late time points on successful and failed Stop trials with differences seeming to account for stopping success. Thus, stop-related changes in SICI and CSP duration seemed more aligned with the Pause and Cancel processes, respectively. The present work provides further evidence from motor system physiology that action-stopping is influenced by multiple inhibitory control processes. Future work should aim to identify activity in cortico-subcortical circuits that could influence motor activity to achieve action-stopping.

## Acknowledgements

We thank Josephine Amick and Tarush Bhatia for assisting with some of the data collection in Experiment 1.

## Author Contributions

Joshua R. Tatz: Conceptualization, Methodology, Software, Formal analysis, Investigation, Visualization, Writing – Original draft preparation, Writing – Reviewing & editing, Supervision; Project administration. Madeline O. Carlson: Investigation, Formal analysis. Carson Lovig: Investigation, Formal Analysis. Jan R. Wessel: Conceptualization, Methodology, Writing – review and editing, Supervision, Project administration, Funding acquisition.

